# Selfish mutations dysregulating RAS-MAPK signaling are pervasive in aged human testes

**DOI:** 10.1101/314815

**Authors:** Geoffrey J. Maher, Hannah K. Ralph, Zhihao Ding, Nils Koelling, Hana Mlcochova, Eleni Giannoulatou, Pawan Dhami, Dirk S. Paul, Stefan H. Stricker, Stephan Beck, Gilean McVean, Andrew OM Wilkie, Anne Goriely

## Abstract

Mosaic mutations present in the germline have important implications for reproductive risk and disease transmission. We previously demonstrated a phenomenon occurring in the male germline, whereby specific mutations arising spontaneously in stem cells (spermatogonia) lead to clonal expansion, resulting in elevated mutation levels in sperm over time. This process, termed *selfish spermatogonial selection*, explains the high spontaneous birth prevalence and strong paternal age-effect of disorders such as achondroplasia, Apert, Noonan and Costello syndromes, with direct experimental evidence currently available for specific positions of six genes (*FGFR2*, *FGFR3*, *RET*, *PTPN11*, *HRAS* and *KRAS*). We present a discovery screen to identify novel mutations and genes showing evidence of positive selection in the male germline, by performing massively parallel simplex PCR using RainDance technology to interrogate mutational hotspots in 67 genes (51.5 kb in total) in 276 biopsies of testes from 5 men (median age: 83 years). Following ultra-deep sequencing (~16,000x), development of a low-frequency variant prioritization strategy and targeted validation, we identified 61 distinct variants present at frequencies as low as 0.06%, including 54 variants not previously directly associated with selfish selection. The majority (80%) of variants identified have previously been implicated in developmental disorders and/or oncogenesis and include mutations in six newly associated genes (*BRAF, CBL, MAP2K1*, *MAP2K2, RAF1* and *SOS1*), all of which encode components of RAS-MAPK pathway and activate signaling. Our findings extend the link between mutations dysregulating the RAS-MAPK pathway and selfish selection, and show that the ageing male germline is a repository for such deleterious mutations.

## INTRODUCTION

The timing, location and functional effects of spontaneous mutations determine the distribution and phenotypes of mutant cells within the body: this can have a variety of impacts on the health of an individual, and potentially, their offspring. Spontaneous mutations occurring during early post-zygotic development lead to widespread tissue mosaicism that, depending on context, may be phenotypically undetectable or cause so-called ‘somatic’ disorders (Campbell et al. 2015). Such early post-zygotic mosaicism occurs commonly, with up to 22% of apparently *de novo* point mutations (DNMs) detectable in a child’s blood sample likely to have occurred after fertilization (Acuna-Hidalgo et al. 2015; Krupp et al. 2017). A corollary is that a further ~4-10% of DNMs and ~4% of copy-number variants (CNVs) present in a child can be detected at low-level in one of the parent’s somatic (usually blood or saliva) samples, and are therefore in fact inherited; as these would have occurred early during parental post-zygotic development (before the separation of the somatic and gonadal lineages), they are associated with a significant risk of recurrence (Campbell et al. 2014; Acuna-Hidalgo et al. 2015; Rahbari et al. 2016; Krupp et al. 2017). By contrast, spontaneous mutations occurring postnatally contribute to tissue-specific, low-level mosaicism, formation of benign tumors, or cancer, depending on the functional consequence(s) of the acquired mutation(s), the clonal dynamics of the tissue involved and the state of the niche (Klein et al. 2010a; Vermeulen et al. 2013; Holstege et al. 2014; Swanton 2015). This latter phenomenon has been documented in apparently healthy somatic tissues that display stem cell replacement (e.g. skin, colon, small intestine and blood), where low levels (~1-10%) of clonal mutations are prevalent and their incidence and frequency increase with age (Hafner et al. 2010; Laurie et al. 2012; Genovese et al. 2014; Jaiswal et al. 2014; Martincorena et al. 2015; McKerrell et al. 2015; Acuna-Hidalgo et al. 2017; Coombs et al. 2017; Martincorena et al. 2017; Zink et al. 2017).

Analogous to the postnatal occurrence of somatic mutations, we previously demonstrated a similar phenomenon, termed selfish spermatogonial selection, that occurs in the testes of adult men as they age. However, because the testis contains germ cells that, upon fertilization, will carry the genetic information across generations, this process has important reproductive implications, being associated with an increased prevalence of pathogenic DNMs in the next-generation. Despite the relatively low average human germline point mutation rate of ~1.2 x 10^-8^ per nucleotide per generation (Kong et al. 2012; Goldmann et al. 2016; Jonsson et al. 2017), specific ‘selfish’ DNMs in *FGFR2*, *FGFR3*, *HRAS*, *PTPN11* and *RET* are observed up to 1000-fold more frequently in offspring (Goriely and Wilkie 2012). These pathogenic mutations, which cause developmental disorders that show an extreme paternal bias in origin and an epidemiological <u>p</u>aternal <u>a</u>ge <u>e</u>ffect (collectively referred to as PAE disorders; for example achondroplasia, Apert, Costello and Noonan syndromes, multiple endocrine neoplasia type 2a/b), are identical (or allelic) to oncogenic driver mutations in tumors (Goriely and Wilkie 2012). We proposed that although the mutational events arise at low background rates in male germ cells, selfish mutations confer a selective advantage to spermatogonia leading to their clonal expansion, which results in increased apparent mutation levels in sperm over time (Goriely and Wilkie 2012; Maher et al. 2014).

Three methods have previously been used to detect selfish mutations in the male germline, each of which has been limited in their ability to evaluate the process at scale: (1) quantification in sperm, (2) quantification in testis biopsies and (3) direct identification in seminiferous tubules. Detecting selfish mutations in sperm, in which individual mutations are present at levels ranging from 10^-3^ to <10^-6^, requires ultra-sensitive techniques that have limited quantitative analysis to small regions of 1-6 nucleotides across five locations in *FGFR2* (x2) (Goriely et al. 2003; Goriely et al. 2005; Yoon et al. 2009), *FGFR3* (x2) (Tiemann-Boege et al. 2002; Goriely et al. 2009) and *HRAS* (Giannoulatou et al. 2013) (Supplementary Table 1). To circumvent the technical challenges caused by mutational dilution within an entire ejaculate, mutations may alternatively be identified following systematic dissection and sequencing of DNA extracted from discrete testicular biopsies. The germ cells (from diploid spermatogonia to haploid spermatozoa) are located in long (up to ~80 cm) highly convoluted and tightly packed seminiferous tubules, comprising ~300-500 per testis (Glass 2005). As clonally expanding mutant spermatogonia are physically restricted to the tubules in which they arise, their geographical distribution within the testis is confined to specific regions: the existence of such localized foci has been demonstrated for selfish mutations in four genes (*FGFR2*, *FGFR3*, *PTPN11*, *RET*) (Qin et al. 2007; Choi et al. 2008; Dakouane Giudicelli et al. 2008; Choi et al. 2012; Shinde et al. 2013; Yoon et al. 2013; Eboreime et al. 2016). Finally, mutant clones have been directly visualized in sections of formalin-fixed paraffin embedded (FFPE) normal human testes using immunohistochemical approaches to reveal abnormal expression of spermatogonial antigens (Lim et al. 2012; Maher et al. 2016a). Microdissection of tubules exhibiting enhanced antigen staining and subsequent whole genome amplification facilitated screening of over 100 genes, identifying 9 new selfish mutations, including one in a novel gene (*KRAS*) (Supplementary Table 1). However this approach is limited both by the need to source fixed testis samples with good tissue morphology and DNA preservation, and by the high threshold required for successful immunohistochemical detection (Maher et al. 2016a; Maher et al. 2016b).

Owing to the limitations outlined above, experimental evidence of clonal expansion has so far been restricted to activating mutations at 16 codons in only six genes (Supplementary Table 1), all encoding members of the receptor tyrosine kinase (RTK)-RAS-MAPK signaling pathway. Here, we hypothesized that other variants dysregulating the RAS-MAPK pathway, and/or other pathways controlling spermatogonial stem cell homeostasis, may be under positive selection in the male germline (Goriely and Wilkie 2012; Goriely et al. 2013). To reduce the required assay sensitivity compared with bulk semen analysis, and hence substantially widen the extent of the genomic target that could feasibly be analyzed in a single experiment, we exploited approach (2) above. By combining systematic dissection of 276 testicular biopsies from 5 individuals with massively parallel simplex PCR and ultra-deep sequencing (~16,000x) of mutational hotspots in 67 genes, we present the most comprehensive survey of mutations clonally enriched in the human testis to date. We describe the identification of 61 distinct variants across 15 genes with variant allele frequencies (VAF) as low as 0.06%, including 51 mutations and 6 novel genes with strong support for association with the process of selfish spermatogonial selection.

## RESULTS

To perform a discovery screen and identify novel mutations and genes under selection in the male germline, we systematically biopsied human testes following the experimental design summarized in Supplementary Figure 1. A total of 276 small biopsies (~60–180 mm^3^) from 5 men (age range 34-90 years, median 83 years) were screened by ultra-deep Illumina sequencing (~16,000x post-filtering) of a panel of candidate loci (corresponding to 66.5 kb genomic sequence across 500 amplicons, covering mutational hotspots in 71 genes; see Methods for criteria used to include loci in screen), amplified using massively parallel simplex PCR (RainDance Thunderstorm). To detect low level mosaicism (~0.1-3.0%), the background at each genomic location was independently estimated for all 431 (of 500) amplicons (in 67 of 71 genes) that passed quality control (Supplementary Table 2). After normalization, a statistical model was applied to call outlier non-consensus variants at each genomic position (within each amplicon): a minimum threshold of 10 variant reads and median coverage of > 5,000x was implemented to reduce false positive calls. As a conservative prioritization strategy, only variants with two or more independent calls were further studied, resulting in a set of 374 variant calls located at 361 genomic locations (see Methods). Visualization and manual curation of each of these calls identified 115 higher confidence candidate variants, distributed at 105 genomic positions across 165 biopsies (Supplementary Figure 1 and Supplementary Table 3).

As calling variants at low levels (<1%) is subject to PCR artefacts and sequencing errors (Minoche et al. 2011; Hestand et al. 2016; Salk et al. 2018), we developed a tiered strategy for further variant prioritization. We reasoned that variants called independently in overlapping amplicons or in sample replicates (12 biopsies were amplified and sequenced in duplicate) were least likely to be artefactual (Tier 1 variants, Table 1). 18 of the 40 Tier 1 variants (with VAF ranging from 0.10% to 2.63%) were re-screened by PCR or using single molecule molecular inversion probes (smMIPs) and ultra-deep MiSeq sequencing (~30,000x). Seventeen of the 18 (94%) variants were validated, suggesting the great majority of Tier 1 variants are true positive calls (Table 1, Supplementary Table 3). Amongst the Tier 1 variants are five mutations previously associated experimentally with selfish selection: *FGFR2* c.755C>G (p.Ser252Trp – Apert syndrome), c.758C>G (p.Pro253Arg – Apert syndrome) and c.870G>T (p.Trp290Cys – Pfeiffer syndrome), *KRAS* c.182A>G (p.Gln61Arg – oncogenic) and *PTPN11* c.215C>T (p.Ala72Val – oncogenic) (Table 1). This strong enrichment for canonical examples of selfish mutations (Supplementary Table 1) provided initial validation of our experimental approach and starting hypothesis.

**Table 1.**
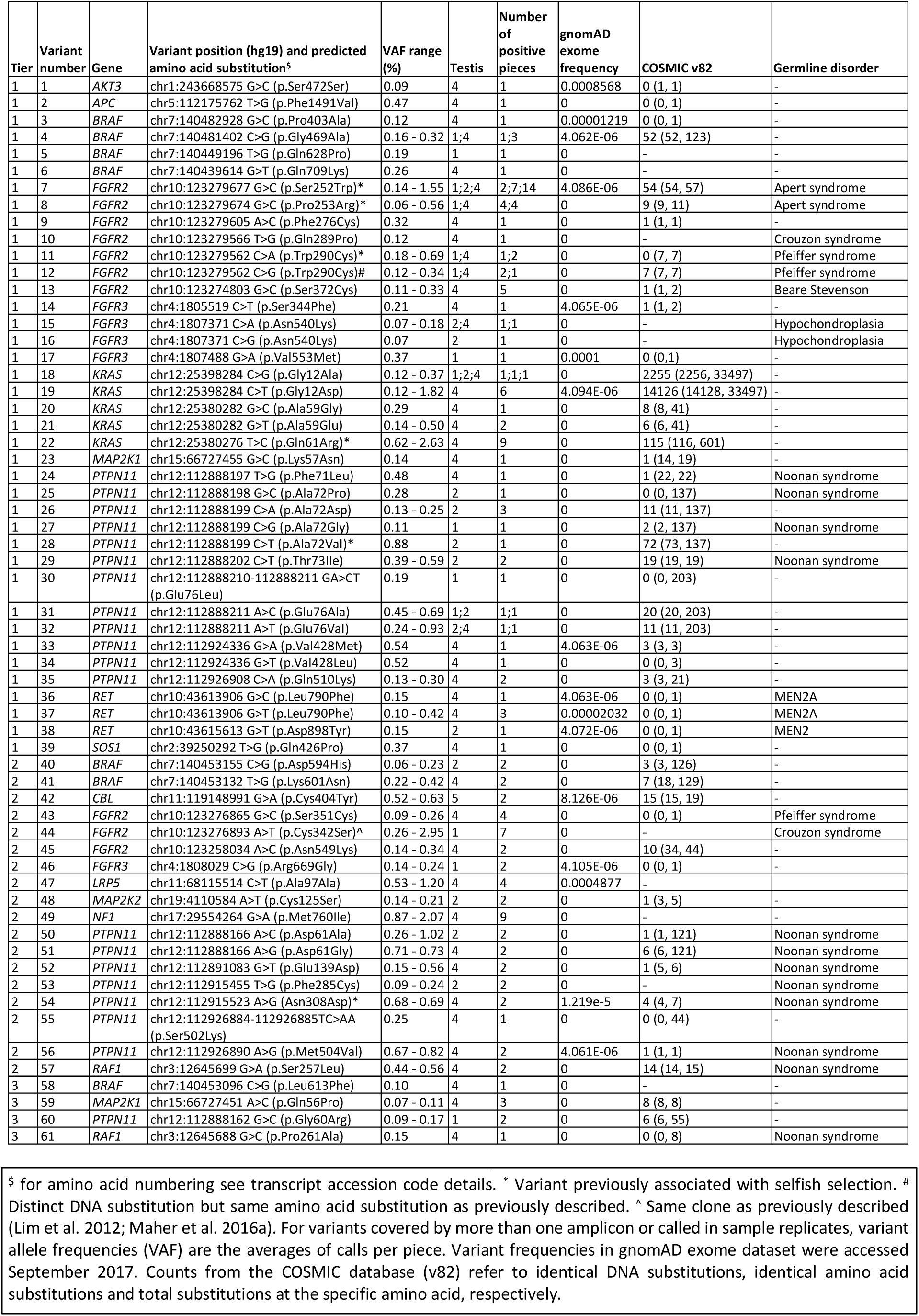
List of 61 validated variants identified in this study.

Within the panel, the majority (88.7%) of callable (i.e. excluding primer sequences and amplicons with low QC) regions were represented by a single amplicon and only 12 biopsies were sequenced in duplicate (Supplementary Table 4): hence, we next investigated variants that were called in single amplicons in two or more biopsies, at VAF of ≥0.2% in at least one biopsy (Tier 2). Twenty-six Tier 2 variants were identified, 18 (69%) of which were validated upon resequencing (Table 1, Supplementary Table 3). Notably, all (14/14) of the known pathogenic variants were validated, but only four of the twelve variants without prior disease association were true positives. In biopsy 4D25, *PTPN11* c.1504T>A (p.Ser502Thr - Noonan syndrome) was called as a single nucleotide variant but on validation it was identified as a double nucleotide substitution c.1504_1505delTCinsAA (p.Ser502Lys). Next, 29 variants with a VAF of 0.1 - <0.2% called in a single amplicon in two or more biopsies (Tier 3) were identified. Only 4 of the 22 (18%) resequenced Tier 3 variants were validated, suggesting that in this lower frequency range, the majority of calls are artefactual (Table 1, Supplementary Table 3). Owing to the low validation rate of variants with VAFs of 0.1 - <0.2%, none of the remaining 20 calls that exhibited VAF <0.1% (Tier 4) variants were re-screened for validation (Supplementary Table 3).

Overall we identified 61 distinct variants that we classified as independently validated, present in 15 of the 67 genes that passed quality control and were analyzed in the experiment. Based on the identification of the same variant in testes sourced from different men, we conclude that at least 72 independent mutational events (clones) could be distinguished across the five testes (Table 1, Figure 1, Supplementary Figs 2-3). Two variants (*FGFR2* c.755C>G (p.Ser252Trp) (#7) and *KRAS* c.35G>C (p.Gly12Ala) (#19)) occurred in three testes and seven in two testes (Figure 1; Supplementary Fig 2). Strikingly, these variants are all either recurrent mutations causative of congenital skeletal disorders, or known hotspots in cancer (COSMIC) that may be associated with lethal or as yet undescribed congenital disorders (Table 1). Figure 2 details all validated variants for the two genes most highly represented in this list, *FGFR2* and *PTPN11* (15 independent mutational events responsible for 10 distinct variants in *FGFR2* (encoding nine pathogenic protein changes); and 22 independent mutational events of 20 distinct variants in *PTPN11*). Their relative locations on the respective protein products shows striking overlap with mutational hotspots previously associated with developmental disorders and cancer. The corollary is that our observations of these mutations in testes are likely to be relevant to the biological origins of the cognate diseases. Similar plots for 13 other genes with validated variants are presented in Supplementary Figure 3.

**Figure 1.**
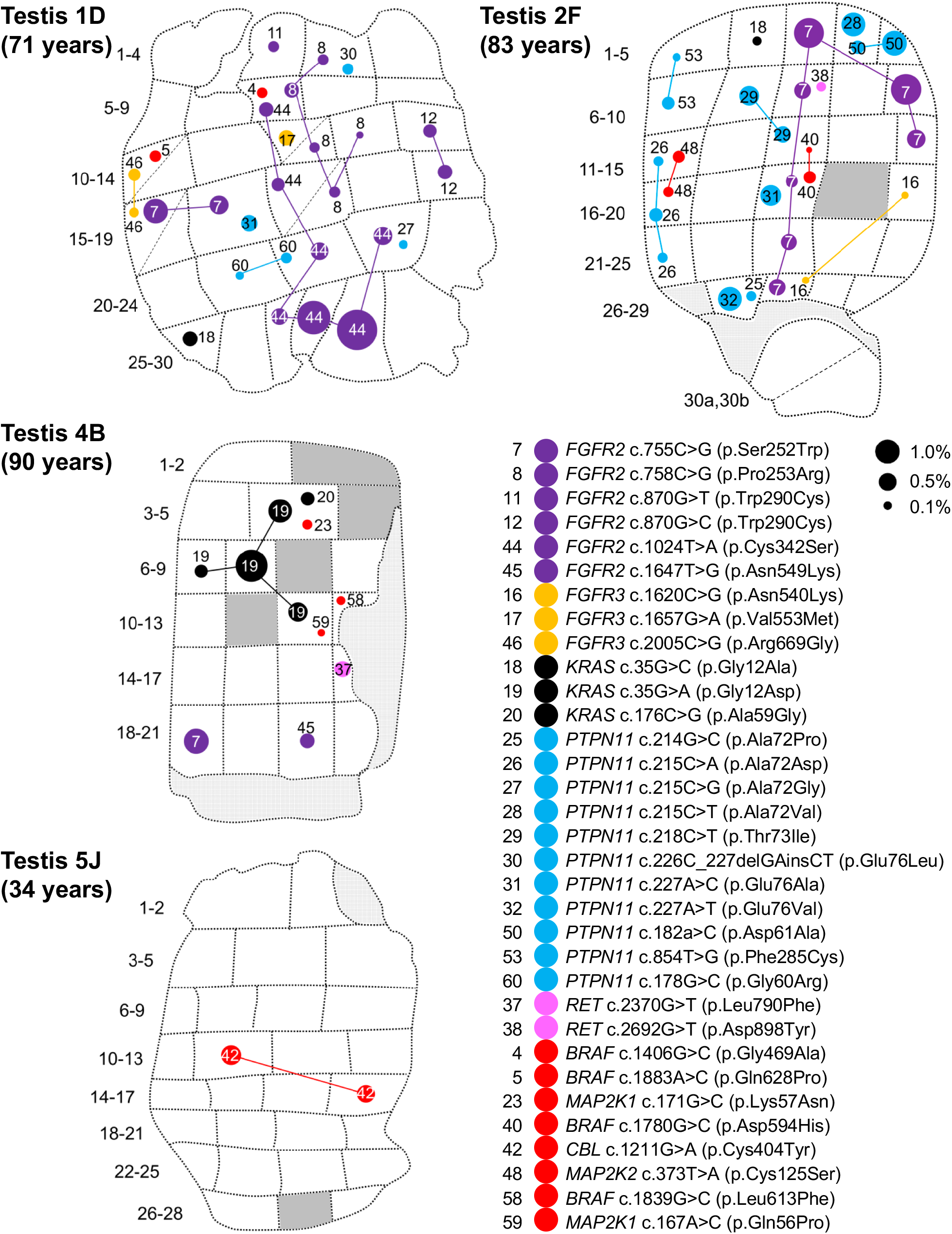
Distribution of validated variants in testis slices 1D, 2F, 4B, and 5J. Testicular biopsy numbers are located to the left of each testis slice. Some biopsies were further dissected into two pieces of which the orientation is unknown – these are indicated with a diagonal dashed line (e.g. Tes2F 30a,b). Each variant has a distinct number (as listed in Table 1) and is colored according to gene: *FGFR2* (purple), *FGFR3* (orange), *KRAS* (black), *PTPN11* (blue), *RET* (pink), newly associated gene (red). The size of each circle is proportional to the observed variant allele frequency (VAF) in each biopsy as indicated by black dots on the figure key. Identical variants in different biopsies have been connected by lines that likely track the seminiferous trajectory across the testis and therefore may represent a single ‘clonal event’; note that the path of the clone has been arbitrarily drawn and may not represent the true geography of a seminiferous tubule across the testis. Dark gray segments represent biopsies that were not sequenced due to insufficient material quality/quantity (see Methods). Light gray segments represent non-tubular regions of tissue. The age of the individual from whom the sample was collected is indicated on the figure (See Supplementary Table 5 for further details on the testicular samples). The remaining five slices of Tes4 are presented in Supplementary Figure 2. Tes3D is omitted as no variants were identified

**Figure 2.**
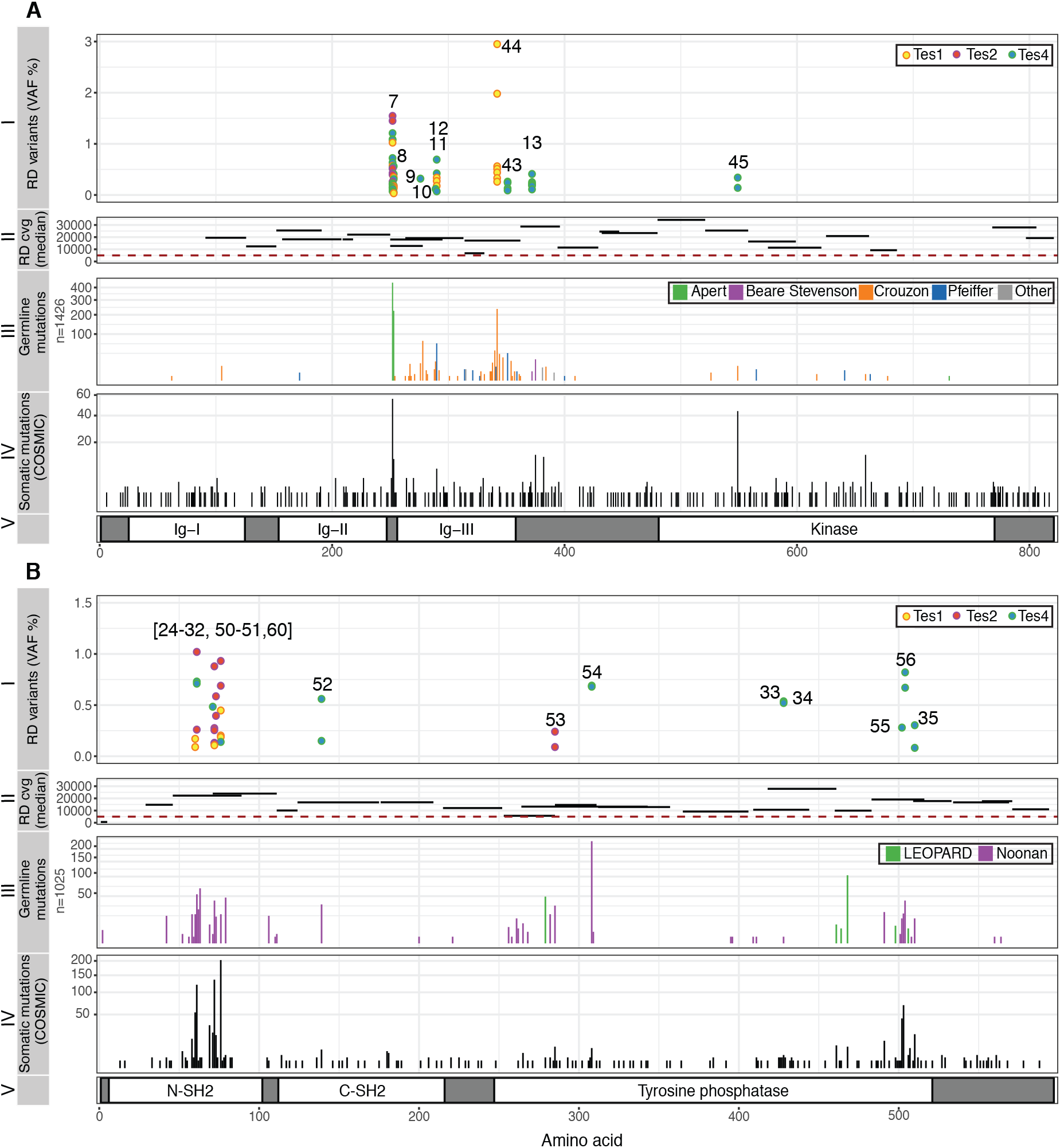
Spontaneous mutations in *FGFR2* (A) and *PTPN11* (encoding SHP2) (B) identified in testicular biopsies. (A) (I) Ten validated variants positioned along the amino acid sequence of FGFR2 (*x*-axis, see panel V), ranging in VAF from 0.06% to 2.95% (*y*-axis), identified in Tes1D, Tes2F and Tes4. Numbers correspond to those in Table 1; two different variants (c.870G>C or T) predicted to cause the same p.Trp290Cys substitution (#11, #12) were identified. (II) Relative location and length of amplicons used to sequence main hotspots of *FGFR2* are plotted on the *x*-axis. Median coverage per amplicon is plotted on the *y*-axis. All amplicons had median coverage above the cut-off (red dashed line) of 5,000x. (III) Number of reported constitutional variants encoding amino acid substitutions in FGFR2 associated with developmental disorders (sqrt scale) (updated from (Wilkie 2005)). (IV) Number of reported somatic amino acid substitutions in FGFR2 in cancer (COSMIC v82). (v) Protein domains of FGFR2. Annotations and protein structure are based on transcript ID NM_000141 and Uniprot ID P21802 (v2017_01), respectively. (B) (I) Twenty validated variants positioned along the amino acid sequence of SHP2 (*x*-axis, see panel (V), ranging in VAF from 0.09% to 1.02% (*y*-axis), identified in Tes1D, Tes2F and Tes4. (II) Location and size of amplicons used to sequence main hotspots of *PTPN11* are plotted on the *x*-axis. Median coverage per amplicon is plotted on the *y*-axis. All amplicons except one had median coverage above the cut-off of 5,000x. (III) Number of reported constitutional variants encoding amino acid substitutions in SHP2 associated with developmental disorders (sqrt scale). (IV) Number of reported somatic amino acid substitutions in SHP2 in cancer (COSMIC v82). (V) Protein domains of SHP2. Annotations and protein structure are based on transcript ID NM_002834 and Uniprot ID Q06124 (v2017_01), respectively.

Next, using the geographical register of the multiple biopsies, the spatial distribution of each variant across the testicular biopsies was investigated (Figure 1, Supplementary Figure 2). For example, in 6 of 153 biopsies across three slices from Tes4 we identified a *KRAS* c.35G>A (p.Gly12Asp) mutation (#18). *KRAS* c.35G>A is one of the most frequently reported substitutions in cancer (>14,000 records in COSMIC v82) and post-zygotic *KRAS* c.35G>A mutations have been reported to cause arteriovenous malformations of the brain (Nikolaev et al. 2018) and linear nevus sebaceous syndrome (Wang et al. 2015), but it has never been reported as a constitutional mutation. In slice 4B (slice B of Testis 4) (Figure 1 and Supplementary Figure 3), this *KRAS* mutation was detected at VAF ranging from 0.26% to 1.82% in four adjacent biopsies, suggestive of an expansion of a mutational event tracking along the length of a single seminiferous tubule. The same *KRAS* variant was also detected in two neighboring biopsies from slices 4D and 4E, apparently at a distance from the larger clone in slice 4B (Supplementary Figure 2); this smaller clone may represent a distinct mutational event having occurred in an independent tubule, but the resolving power of the experiment does not exclude the possibility that this is a large clonal event spreading along the length of a single seminiferous tubule (that measure up to ~80 cm in humans).

Owing to the convoluted packing of the seminiferous tubules, individual testicular biopsies contain segments of multiple individual tubules and in 43 biopsies more than one variant was identified (Figure 1, Supplementary Figure 2 and Supplementary Table 3). Mutations with similar distributions across multiple biopsies may represent clones either within the same tubule, or in distinct intermingled tubules running alongside each other. For example, two distinct mutations, *MAP2K2* c.373T>A (p.Cys125Ser) (oncogenic) and *PTPN11* c.215C>A (p.Ala72Asp) (oncogenic)] are both found in the adjacent biopsies 2F11 and 2F16 (Figure 1), with the latter mutation extending into the neighboring biopsy 2F21. In Tes4, four of the six biopsies positive for the oncogenic *KRAS* c.182A>G (p.Gln61Arg) mutation (4E18, 4E25, 4F27, 4G1) were also positive for a synonymous variant in *LRP5* [c.291C>T (p.Ala97Ala); no prior disease association] (Supplementary Figures 2 and 4).

In contrast to selfish mutations that arise in adult spermatogonia and are therefore restricted to the seminiferous tubules in which they arise, ‘classical’ post-zygotic mosaic mutations occurring in embryonic primordial germ cells, before the formation of the seminiferous tubules, are expected to have a wider distribution in one or both testes. We found one suggestive example of this, an *NF1* c.2280G>A (p.Met760Ile) variant, which exhibited a pattern of occurrence in Tes4 distinct from all the other identified mutations. The variant was originally called in nine biopsies at relatively high VAF (median 1.1%, range 0.9-2.1%) (Supplementary Figure 2), and inspection of the mutation frequency in each sample (Supplementary Figure 5) showed numerous other biopsies in Tes4 with elevated VAFs, compatible with an earlier post-zygotic mosaic event. Unfortunately, no other tissue was available from this individual to test whether the variant was restricted to a single testis and/or to the germline tissue.

To explore the relationship between mutational events identified using RainDance technology (which inherently involves destruction of the tissue structure of the testis) and the occurrence of mutations in individual seminiferous tubules, we exploited the availability of adjacent FFPE material for two of the testes. In Tes1D, our deep-screening strategy identified a *FGFR2* c.1024T>A (p.Cys342Ser) variant at VAFs ranging from 0.26% to 2.95% in seven contiguous biopsies, suggestive of a clonal event tracking a single seminiferous tubule across the testis (Figures 1 and 2, variant #44). For this testis, we had previously studied the adjacent FFPE tissue block (Tes1-1 described in (Lim et al. 2012; Maher et al. 2016a)) using immunohistochemical staining for markers of selfish clones (enhanced MAGEA4 and pAKT immunostaining), followed by laser capture microdissection and targeted resequencing.

Strikingly, we previously identified and validated the identical *FGFR2* variant, strongly suggesting that this large mutant clone is present within a significant portion of a single seminiferous tubule that tracks across adjacent testis slices (Maher et al. 2016a). To seek further examples, we undertook a new analysis of putative mutant clones within Tes2E, a FFPE tissue block adjacent to the Tes2F slice, to identify individual tubular cross-sections exhibiting enhanced MAGEA4 immunostaining; laser capture microdissection of six distinct groups of tubular cross-sections, followed by PCR and Illumina sequencing confirmed the presence of the *FGFR2* c.755C>G (p.Ser252Trp – Apert syndrome) and *PTPN11* c.214G>C (p.Ala72Pro – Noonan syndrome) mutations in distinct enhanced MAGEA4-tubules, consistent with the geographic location of these specific variants identified by deep-sequencing in the adjacent Tes2F slice (Figure 3). For the three other testes, FFPE blocks were not available.

**Figure 3.**
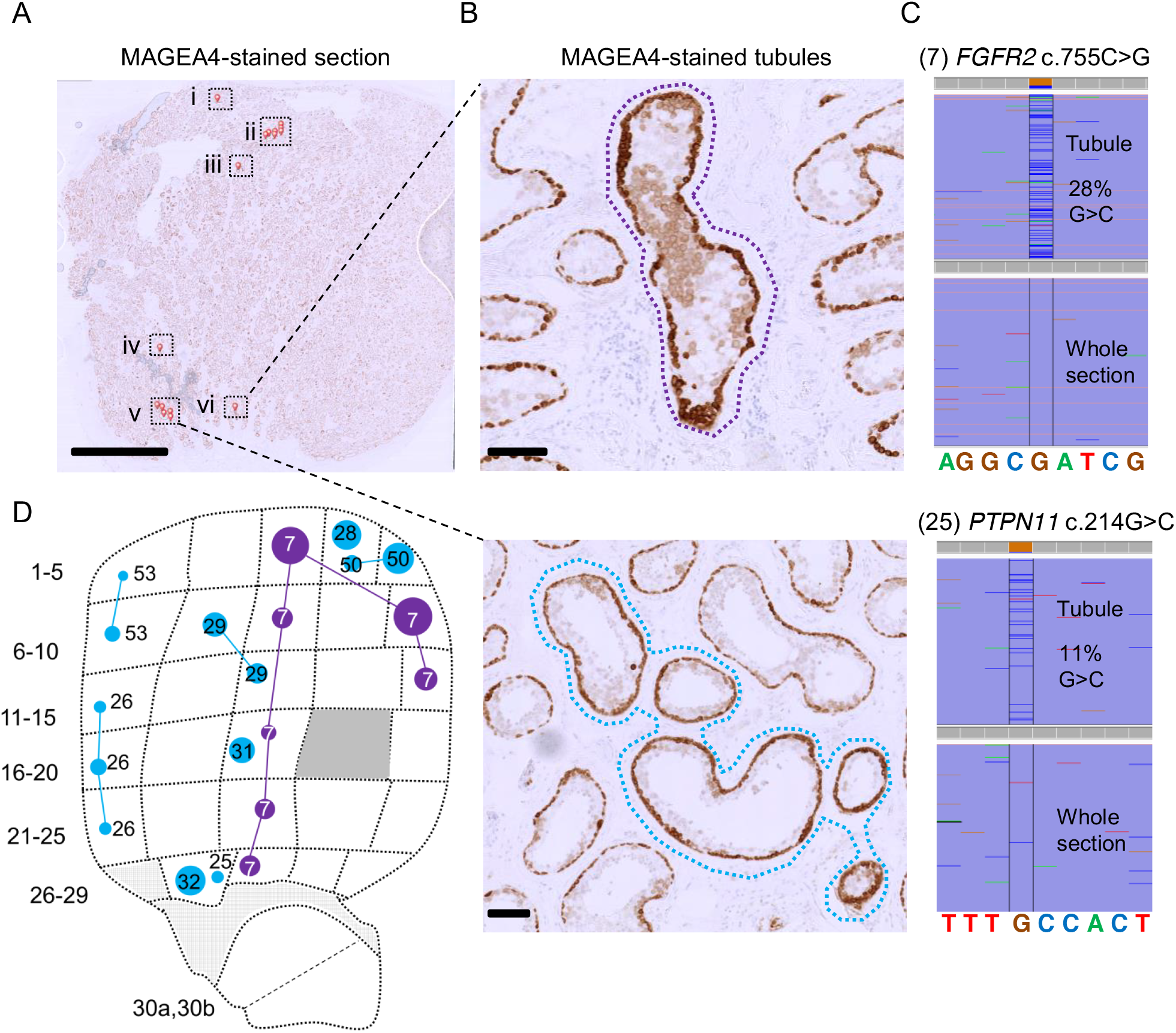
Visualization of mutant tubules in Testis 2. (A) A 5 μm thin section from Tes2E, a FFPE block of tissue adjacent to the testis slice 2F, immunostained with anti-MAGEA4 antibody to label spermatogonia. Seminiferous tubules with enhanced MAGEA4 immunopositivity, suggestive of the presence of mutant clones are labelled with small red pins and boxed. Scale bar = 5 mm. (B) High magnification view of cross-sections with MAGEA4-enhanced immunopositivity in two localized areas are labelled with dotted lassoes representing the laser-microdissected regions. Scale bars = 100 μm. (C) Results from targeted resequencing of the microdissected seminiferous tubules labelled by dotted lassoes in (B) viewed in IGV (Integrated Genome Viewer); spontaneous pathogenic *FGFR2* c.755C>G #7 (top) and *PTPN11* c.214G>C #25 (bottom) variants were identified in DNA extracted from microdissected tubule cross-sections, but not in DNA from the whole tissue section. Comparison of the MAGEA4 section (A) with adjacent testis slice 2F from the Raindance screen (D) (the same image as in Figure 1 but showing only the targeted *FGFR2* and *PTPN11* mutations), shows that both variants match to a mutation previously identified in the corresponding position of testis slice 2F.

## DISCUSSION

We present a new broad-scale approach to studying clonal *de novo* germline mutations directly in human adult testes, the tissue where the majority of DNMs originate. Utilizing massively parallel multiplex PCR and ultra-deep sequencing of 51.5 kb in 276 discrete human testicular biopsies followed by the implementation of a statistical prioritization calling strategy, we identified 61 different variants in a total of 111 mutation-positive biopsies, 59 of which encode non-synonymous substitutions (Table 1).

Several observations support the notion that the mutations identified are strongly enriched for clonal events that are promoted by positive selection of mutant stem cells via the phenomenon of selfish spermatogonial selection. Out of the 61 validated variants (Table 1), 43 are located in five (*FGFR2*, *FGFR3*, *KRAS*, *PTPN11*, *RET*) of the six genes associated with strong prior experimental evidence for this process (Supplementary Table 1). As detailed in Table 1 and illustrated in Figure 2 and Supplementary Figure 3, the vast majority of variants identified across these five genes overlap with those observed in dominant congenital disorders and/or cancer, strongly suggestive of a functional role via a gain-of-function mechanism. The most commonly observed individual mutation was *FGFR2* c.755C>G (p.Ser252Trp - Apert syndrome) detected in 23 biopsies. In this and other cases, the identification of identical variants in multiple neighboring testis biopsies (Figure 1 and Supplementary Figure 2) is supportive of clonal expansion along the length of the seminiferous tubules, and in three cases this process could be directly validated at a cellular level by visualizing the selfish expansion characterized by enhanced MAGEA4 staining in the adjacent testis block (Figure 3 and (Maher et al. 2016a)). The largest number of mutations was observed for *PTPN11* (encoding the SHP2 tyrosine phosphatase), in which we identified 20 different variants (across 33 biopsies) (Table 1 and Figure 2B). We observed 12 distinct variants located within the N-SH2 domain of SHP2, a region of the protein known to repress the catalytic phosphatase domain in its wild-type state (Neel et al. 2003), including each of the possible nucleotide substitutions at *PTPN11* c.215C encoding three distinct amino acids (p.Ala72Asp, p.Ala72Gly and p.Ala72Val) that have been associated with Noonan syndrome or oncogenesis. This wide mutational spectrum is consistent with epidemiological data that concur that *PTPN11*-associated Noonan syndrome mutations have a high spontaneous birth prevalence (~1/10,000 births) (Goriely and Wilkie 2012). We also identified two dinucleotide substitutions in *PTPN11*: both the c.226_227delGAinsCT (p.Glu76Leu (#30)) and the c.1504_1505delTCinsAA (p.Ser502Lys (#55)) variants encode amino acid substitutions that, owing to the nature of the genetic code, cannot arise from single-nucleotide changes. These observations are reminiscent of other previously described selfish mutations encoded by double and triple substitutions, which in some cases, were shown to result via a ‘double-hit’ mechanism (Goriely et al. 2005; Goriely and Wilkie 2012; Giannoulatou et al. 2013). In humans, the *de novo* tandem mutation rate is estimated to be ~0.3% of the single nucleotide variant rate (Besenbacher et al. 2016); in this small set of 61 variants, we find a ~10-fold enrichment over the background rate.

Given this strong support for positive clonal selection of pathogenic variants in previously known selfish genes, the next question is whether the other 18 validated variants present in novel candidate genes might also signal the presence of selfish selection. We first excluded from consideration one variant, *NF1* c.2280G>A p.(Met760Ile) (variant #49), which presented with a different pattern of occurrence characterized by an extended geographical distribution across ~1/3 of the testis from individual Tes4, raising the possibility of an early post-zygotic (as opposed to adult-onset) mutational event (Supplementary Figure 5). Although this NF1 variant exhibits a high CADD (24.6)/Polyphen score, has been reported in one case of lung cancer (Redig et al. 2016) and is located within the cysteine-serine-rich domain (CSRD), a region where several missense mutations associated with breast cancer and neurofibromatosis have been identified (Koczkowska et al. 2018), its pathogenic status - and potential for positive selection - remain uncertain.

Of the remaining 17 variants, all but three are accounted for by six genes (*BRAF*, *CBL*, *MAP2K1*, *MAP2K2*, *RAF1* and *SOS1*) encoding members of the RAS-MAPK pathway, among which nine variants have previously been reported in either congenital disorders or cancer (Table 1 and Supplementary Figure 3). Moreover, for several variants (BRAF p.Gly469Ala, MAP2K1 p.Lys57Asn and p.Gln56Pro, MAP2K2 p.Cys125Ser, RAF1 p.Ser257Leu and p.Pro261Ala), direct biochemical evidence of a dominant gain-of-function activity is available (Wan et al. 2004; Kobayashi et al. 2010; Van Allen et al. 2014; Arcila et al. 2015). In fact, only three validated variants (#1,2,47), for which evidence of involvement in selfish selection is weak or can be ruled out, were found in genes (*APC*, *AKT3*, *LRP5*) that function outside the RTK-RAS-MAPK pathway (see Supplementary Note). Hence, although only 41.9% of the callable sequence of our panel comprised RTK-RAS-MAPK candidate genes, 95% (57/60) of the validated variants represented known or very likely pathogenic changes within members of this signaling pathway (p value = 4.233e-13, Fisher’s two tailed test), reinforcing the proposal that activation of the RAS-MAPK pathway is the predominant mechanism underlying selfish spermatogonial selection (Goriely et al. 2003; Goriely et al. 2009; Goriely and Wilkie 2012; Maher et al. 2016a). Mutations in other core cellular pathways in human testes may either not be associated with positive selection or may lead to milder clonal expansions that will require more sensitive screening approaches to uncover. Although it can be difficult formally to distinguish signals of selection from normal turnover/neutral drift dynamics whereby the random loss of some clones is compensated by the expansion of others over time (Klein et al. 2010b; Simons 2016; Zink et al. 2017), the highly significant enrichment of functionally significant (biochemically activating) mutations affecting a single signaling pathway argues against a neutral process.

Among the variants we identified, we observed a high proportion of strongly oncogenic mutations with 23 of the 35 non-synonymous variants reported in COSMIC (v82) having never been described as constitutional mutations (Table 1). Strong gain-of-function mutations would be more likely to promote efficient expansion of spermatogonial stem cells and result in larger clones that are easier to detect. However, in order to be transmitted, the mutations must be compatible with formation of functional sperm and with embryonic development. We previously showed that tubules with spermatogonia harboring strongly oncogenic variants are associated with reduced numbers of post-meiotic cells (Maher et al. 2016a). This would represent a mechanism by which the testis ‘filters’ the transmission of pathogenic mutations across generations, although proof of this concept would require development of ultra-sensitive assays to screen large numbers of sperm samples. It is noteworthy that despite the relative abundance of strongly oncogenic mutations in the adult male germline, testicular tumors originating from adult spermatogonia (spermatocytic tumors) are extremely rare, with an incidence of ^~^1 per million men and are mostly benign in nature (Ghazarian et al. 2015; Giannoulatou et al. 2017).

The age range of the testes analyzed in this study was highly skewed, with four being sampled from older individuals (aged 71-90 years), and one (Tes5J) from a 34-year old man. While for three of the four older individuals we identified multiple mutation-positive biopsies, Tes5J from the younger man contained only two mutation-positive biopsies – likely representing a single clonal event - carrying the oncogenic *CBL* c.1211G>A (p.Cys404Tyr) variant (at VAF 0.5- 0.6%), in keeping with the expectation that the prevalence and size of mutant clones increases with time. It was however surprising that no variants were detected in Tes3D, given the advanced age of the donor (87 years). Although it is possible that this individual may have had a low propensity to accumulation of selfish mutations, a more likely explanation is that few or no germ cells were present in this testis slice, either due to Sertoli-cell only syndrome or due to age-related atrophy (Paniagua et al. 1987). Unfortunately, as no tissue had been preserved for histological analysis, we were unable to determine the status of spermatogenesis in this sample.

Our study has several technical limitations. The majority of variants identified were present at VAFs <1%, close to the typical detection limits attributable to the error rates associated with DNA damage (10^-2^-10^-4^) (Arbeithuber et al. 2016; Chen et al. 2017), PCR (10^-4^-10^-6^) (Hestand et al. 2016; Potapov and Ong 2017) and Illumina sequencing (^~^10^-3^) (Minoche et al. 2011) (Salk et al. 2018). To account for such technical confounders, we employed a conservative custom statistical approach to determine the background error rate at each position and to prioritize variants (Supplementary Figure 1). Although we confirmed variants with a frequency as low as 0.06% using this approach, the majority (81.8%) of the prioritized variants called in single amplicon at VAFs of 0.1-0.2% (Tier 3) were false positives. In the twelve samples amplified and sequenced in duplicate, only 7 of 15 variants were called in both replicates (Supplementary Table 4). The best predictor of true positives was the presence of a call in more than one amplicon (100% validation rate); for calls in single amplicons the best predictor was the pathogenicity of the variant (17 of 18 (94.4%) pathogenic variants vs. 5 of 30 (16.7%) without prior disease association validated). Broad-scale approaches that target both DNA strands and use unique molecular indexes such as Duplex sequencing (Kennedy et al. 2014) or smMIPs (Hiatt et al. 2013) (used here to validate a subset of variants) represent valuable alternatives to direct PCR amplification in future studies to reduce background errors (Salk et al. 2018). Overall 14% of the designed amplicons did not pass quality control (due to insufficient coverage, mapping error…), which included those targeting candidate PAE mutations such as eight mutational hotspots in *FGFR3*, six in *PTPN11*, one in *RET* (p.Val804), and other key hotspots in *SKI* (Shprintzen-Goldberg syndrome), *SETBP1* (Schinzel-Giedion syndrome) and *AKT1* (p.Glu17Lys – Proteus syndrome, oncogenesis). Although considered to be the most frequently mutated nucleotide in the germline with a birth prevalence of ^~^1:30,000 (Bellus et al. 1995), we did not detect the *FGFR3* c.1138G>A or c.1138G>C achondroplasia-associated mutations due to exclusion of this region because of insufficient coverage (<5,000x) (Supplementary Table 2; Supplementary Fig 3E).

In summary this work represents a new approach to studying DNMs directly in their tissue of origin. By utilizing the clonal nature of mutations that leads to focal enrichment, we circumvented the technical difficulties associated with calling DNMs in single sperm or the poor DNA quality associated with immunopositive tubules from FFPE material. In a single biopsy a whole population of *de novo* mutations can be assessed. Studying mutations within the testis facilitates identification of mutations and pathways under positive selection in spermatogonia but that may be incompatible with life, either by impairing gamete differentiation and sperm production or by causing early embryonic lethality. Our approach reveals the prevalence and geographical extent of clonal mutations in normal human testes, suggesting that the ageing male germline is a repository for functionally significant, often deleterious mutations. Based on an estimated total birth prevalence of DNMs causing developmental disorders of 1 in 295 (DDD 2017), such PAE mutations may contribute 5-10% of the total burden of pathological mutations, depending on paternal age. Investigating the clonal nature of spontaneous testicular variants also provides insights into the regulation of the poorly-studied human spermatogonial stem cell dynamics and how spontaneous pathogenic mutations hijack homeostatic regulation in this tissue to increase their likelihood of transmission to the next generation.

## METHODS

### Testis samples

Ethical approval was given for the use of human testicular tissue by the Oxfordshire Research Ethics Committee A (C03.076: Receptor tyrosine kinases and germ cell development: detection of mutations in normal testis, testicular tumors and sperm). Testes from five men aged 34, 71, 83, 87 and 90 years were either commercially sourced or obtained locally from research banks or post-mortems, with appropriate consent (Supplementary Table 5). Each testis was cut into slices ^~^3-5 mm thick and either stored frozen at −80°C or formalin-fixed. After thawing slices of frozen testis, extraneous tissue (epididymis or tunica albuginea) was removed and slices were further dissected into 21-36 biopsies (Supplementary Table 5). Biopsies were pulverized using a pestle and DNA extraction was performed using the Qiagen DNeasy Blood & Tissue Kit. Samples with insufficient DNA quantity (determined using Qubit fluorometer (Life Technologies)) or quality (determined using Nanodrop spectrophotometer (Thermo Scientific)) were excluded, resulting in a total of 276 biopsies [Tes1D (34 biopsies), Tes2F (30 biopsies), Tes3D (32 biopsies), Tes4B-4G (153 biopsies from 6 slices), Tes5J (27 biopsies)].

### RainDance library preparation and sequencing

Primer pairs (tailed with common RainDance sequences (RD)) targeting 500 genomic regions (20-169 bp [average 133 bp, median 143 bp]) in 71 genes (66.5 kb in total) were designed by RainDance Technologies. The panel comprised mutational hotspots in the six established PAE genes, genes encoding other RTKs and members of the RAS-MAPK signaling pathway, genes in other pathways associated with spontaneous disorders that display narrow mutational spectra suggestive of gain-of-function effects but lacking epidemiological data for paternal age-effect, oncogenes commonly mutated in cancer, some of which are also associated with germline disorders, and regions of 10 control genes. Details of all targeted regions and primers used for amplification are provided in Supplementary Table 6. To maximize the number of different molecules amplified, massively parallel simplex PCR was performed using the RainDance Thunderstorm target enrichment system following the manufacturer’s instructions. Briefly, for each sample, 6 μg of genomic DNA (gDNA) was sheared to an average size of 3,000 bp (using a Covaris blue AFA miniTUBE) and purified using a minElute column (Qiagen). One microliter (out of 20 μl) was run on a gel to verify that the gDNA had been sheared to the correct size range and the remaining gDNA was quantified using a Qubit fluorometer (Life Technologies). The custom primer library, 1.75 μg of sheared gDNA and PCR mix (Platinum Taq Polymerase High Fidelity reagents (Invitrogen), 2.5 mM MgSO_4_, 0.35 μM dNTPs, 0.6 M betaine, 7% dimethyl sulfoxide (DMSO), in 25 μl volume) were loaded onto a ThunderStorm enrichment chip (48 samples at a time). Droplets containing up to 5 primer pairs were merged with gDNA droplets to generate an average of 2 x 10^6^ droplets per sample (525,000 haploid genomes; average of 1 haploid genome per 3-4 droplets; ^~^1000 genomes/individual primer pair (Supplementary Figure 1). Following the merge, libraries were PCR-amplified (94°C for 2 min; 54 cycles of 94°C, 54°C, 68°C for 30 s each; 68°C for 10 min) and the emulsion was broken down with 75-100 μl of Droplet Destabilizer (RainDance) before being purified using AMPure beads (Agencourt). An aliquot of each sample was run on a Bioanalyzer high sensitivity chip (Agilent) to verify the amplification profile and determine the sample concentration. Sixteen different Illumina sequencing tailed libraries were constructed using a set of barcoded (8 bp barcode (BC)) Illumina PE2-RD-rev adaptors, a common PE1-RD-Fwd, 4 ng of merged amplicons and Phusion Hot Start Flex DNA Polymerase (New England BioLabs) with 8% DMSO (98°C for 30 s, followed by 10 cycles of 98°C for 15 s, 56°C for 30 s, 72°C for 40 s, and a final extension at 72°C for 10 min). Following purification (Qiagen MinElute), the relative concentration of the secondary tailing PCR samples was estimated by Real-Time PCR using PE1 and PE2 primers. For each of the 16 libraries, 18 samples with BC1-18 were pooled in equimolar ratio and each final library was diluted to 10 nM. A total of 288 samples (264 singletons and 12 in duplicate) were amplified across 6 ThunderStorm enrichment chips (48 samples each) and subsequently ultra-deep sequenced (^~^22,000x) on two flow cells (16 lanes; 18 samples per lane) of Illumina HiSeq 2000 (2 x 100 bp) using RD-Read1 and RD-Read2 custom sequencing primers generating 14-20 x 10^7^ paired-end reads per library.

### Sequence alignment and variant calling and prioritization

Low quality reads with more than 20 bases below Q20, read pairs with one or two short (<50 bp) reads and reads pairs with unmatched or mismatched sequences between the forward and reverse primer pairs expected for each amplicon were removed. Reads passing QC (on average 86% of reads) were aligned to the human genome (hg19) using BWA-MEM version 0.7.10 (Li 2014) with default parameter settings. Primer sequences were included in the alignment but ignored during variant quantification. The Python library Pysam was used to fetch reads mapped to each amplicon and mapped bases (indicated as letter "M") were identified from the CIGAR string. Pileup was then performed for each amplicon independently. Nine amplicons that did not map to the targeted genomic regions were excluded from subsequent analyses (Supplementary Table 2). Reads with more than 10 non-reference bases were removed (<1% of coverage on average). For amplicons shorter than 200 bp, to avoid double-counting reads at positions where Read 1 and Read 2 overlapped, only the base with the higher quality was considered.

Data exploration of the non-consensus variant counts within each amplicon across the different samples revealed clear data structure with differences between flow cells, sequencing lanes, coverage depths and base quality scores. To reduce false-positive calls, primer sequences were trimmed and only variants supported by at least 10 reads were called. To account for the technical confounders, the data were normalized (accounting for flow cell, lane, and average base quality at each position) using a simple linear model

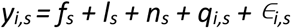

where *y_i,s_* is the nucleotide count for sample *s* at position *i*; *f_s_*, *l_s_* and *n_s_* are the flow cell identifier, the sequencing lane identifier, and individual identifier for sample *s* respectively; and *q_i,s_* is the average base quality of sample *s* at position *i*. We used the glm package in R for model inference (glm(y ^~^ f + l + n + q, family=gaussian())). Values of ∈_*i,s*_ are the normalized signals after accounting for the technical confounders and were used as the inputs for the subsequent analyses. To further account for the effect of the sequencing lane structure, we removed the median effect from each lane to reduce the background signal g_i,s_ = ∈_*i,s*_ – median[∈_*i,s*_ for all *s* in same lane as *s*], before stabilizing the variance using the transformation *g’_i,s_* = *g_i,s_* ∕IQR[*g_i,._*], where IQR[*g_i,._*] is the inter-quantile range at site *i* across all samples. Following these normalization steps, variant calling was performed using a normal model to test for an increase in non-consensus variant calls. Assuming that under the null hypothesis the normalized variant quantification follows a normal distribution 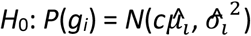 with mean 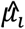 and variance 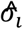, estimated using signals from all samples. We applied a one-sided *z* test in R (pnorm(g, mean=mu0, sd=sigma2, lower.tail=FALSE). Non-consensus calls at each genomic position across the 288 samples were tested independently in each amplicon that passed QC. Variant prioritization was performed using a P-value cutoff of –*log*_10_*P* > 20, which resulted in a total of 19,625 genomic positions with at least one non-reference call.

As samples or amplicons with an excessive number of variants were more likely to represent technical artefacts, these outliers were identified using a Chi Square (*χ^2^*) test, where the expected number of substitutions is defined as the median across all samples. Using a *χ^2^* threshold of –*log*_10_*P* > 3, seven amplicons and 185 sample-mutation combinations were removed from further analysis. Notably, the majority of these were C>A (=G>T) variant calls (Supplementary Figure 4), which represent a known mutational signature associated with oxidative stress that likely arose during sample preparation (Arbeithuber et al. 2016; Chen et al. 2017). Further filtering was performed to remove potential sources of artefacts: calls positioned 1 base from the amplification primer 3’-end were excluded; calls with a maximum VAF of ≥3% were excluded to avoid calling SNPs and to eliminate gross alignment errors or calling of non-consensus variants resulting from homologous genomic regions or pseudogene amplification; positions with a median depth coverage below 5,000x across all samples were excluded (this removed a 53 further amplicons (10.6%) from the analysis; Supplementary Table 2). This resulted in a total of 5729 calls (5659 distinct variants) at 5421 positions, the majority (90.2%) of which were made in a single amplicon and sample. As singleton calls were more likely to represent PCR or sequencing artefacts, we further prioritized calls made in two or more samples and/or present in overlapping amplicons. To exclude potential batch effects, variants were excluded if all calls were made from a single library and the number of calls was >3. This strategy identified 374 variants at 361 genomic positions. VAFs across all samples at each of the 361 genomic positions were plotted and manually inspected for sequencing library preparation or batch effects; raw sequencing reads from calls with suspected sequence misalignment were visualized in Integrative Genomics Viewer (IGV) (Robinson et al. 2011). Variant calls showing evidence of library-specific batch or sequence misalignment effects were excluded from further analysis. Variants in *PTPN11* that matched bases at homologous positions in one of its four pseudogenes were also excluded. The remaining 115 variants at 105 genomic positions were annotated with ANNOVAR version 2015Jun17 (Wang et al. 2010).

### Variant validation

DNA from at least one putative-positive biopsy sample and at least 8 control samples (unrelated blood gDNA and gDNA from other testicular biopsies) was screened by PCR or single molecule molecular inversion probes (smMIPs) (primer and smMIP details in Supplementary Table 6) and sequenced using Illumina MiSeq 300v2 (PCR) or 150v3 (smMIP) kits (further details in Supplementary Methods). Demultiplexed reads were aligned to the human genome (hg19) using BWA-MEM version 0.7.12 (Li 2014). Summary tables of the calls across the aligned target region for PCR and smMIPs were generated using SAMtools mpileup and a custom script (Amplimap – see Supplementary Methods), respectively. A base call was only considered if its mapping quality was ≥Q20 and phred score ≥Q30. Validated variants were annotated according to the following transcripts - *APC*: NM_001127510, *AKT3*: NM_005465, *BRAF*: NM_004333, *CBL*: NM_005188, *FGFR2*: NM_000141, *FGFR3*: NM_000142, *KRAS*: NM_033360, *LRP5:* NM_002335, *MAP2K1*: NM_002755, *MAP2K2*: NM_030662, *NF1*: NM_001042492, *PTPN11*: NM_002834, *RAF1*: NM_002880, *RET*: NM_020975, *SOS1*: NM_005633.

### Immunohistochemistry, microdissection and targeted mutation screen

Where mutations had been identified in frozen sections for which an adjacent FFPE tissue block was available, we attempted to visualize the corresponding mutant clone in sections of the FFPE block. Immunohistochemical staining with anti-MAGEA4 antibody (clone 57B, gifted by Prof. Giulio C. Spagnoli) to identify tubules with enhanced spermatogonial MAGEA4 staining, followed by laser capture microdissection and DNA extraction of adjacent FFPE sections, was performed as described (Maher et al. 2016a). DNA was subsequently amplified by PCR (40 cycles) using CS-tagged primers (Supplementary Table 6) and barcoded for Illumina MiSeq 300v2 sequencing as described above (see also Supplementary Methods). DNA samples extracted from the whole tissue section and from adjacent tubules with a normal MAGEA4 staining appearance were used as controls. Reads were aligned to the human genome (hg19) using BWA-MEM version 0.7.12 (Li 2014) and were visualized in IGV.

### DATA ACCESS

#### Databases and online resources

gnomAD: http://gnomad.broadinstitute.org/

COSMIC: http://cancer.sanger.ac.uk/cosmic/

ClinVar: https://www.ncbi.nlm.nih.gov/clinvar/

OMIM: http://www.omim.org/

## ACKNOWLEDGEMENTS

The authors thank Indira Taylor, Marie Bernkopf and Yan Zhou for technical support, John Frankland and Tim Rostron for dideoxy-sequencing and the High-Throughput Genomics core at the Wellcome Trust Centre for Human Genetics for generation of the Illumina sequencing data. We thank the UCL Cancer Institute Genomics and Genome Engineering Core Facility (supported by the Cancer Research UK – UCL Centre), for providing access to the RainDance Thunderstorm platform, which was purchased on a Wellcome multi-user grant (99148). This work was primarily supported by grants from the Wellcome (grant 091182 to A.G., G.McV. and A.O.M.W.; grant 102731 to A.O.M.W. and studentship 105361 to H.K.R.), the Simons Foundation (332759 to A.G.) and the National Institute for Health Research (NIHR) Oxford Biomedical Research Centre Programme (to A.G.). S.B., P.D. and S.S. were supported by a Wellcome programme grant and D.P. was supported by EU-FP7. We acknowledge funding from the Medical Research Council (MRC) through the WIMM Strategic Alliance (G0902418 and MC_UU_12025) and the support of the High-Throughput Genomics core facility by the Wellcome grant 090532. The funders had no role in study design, data collection and analysis, decision to publish, or preparation of the manuscript.

## Author contributions

Experiments: GJM, HKR, AG; Technical support: HM, PD, DSP, SS, SB; Data analysis: GM, HKR, ZD, NK, EG, GMcV, AG; Manuscript writing: GJM, AOMW, AG; Conception, design and supervision: GMcV, AOMW, AG

## Supplementary material

**Supplementary Figure 1 – Schematic of experimental design.**

**Supplementary Figure 2 – Distribution of mutations in slices Tes4B-4G from individual 4.** Testicular biopsy numbers are located outside and to the left of each testis slice. Each variant has a distinct number (as listed in Table 1) and is colored according to gene: *FGFR2* (purple), *FGFR3* (orange), *KRAS* (black), *PTPN11* (blue), *RET* (pink), newly associated gene (red), NF1 mosaic (yellow with red surround). The size of each circle is proportional to the mutation frequency. Lines connect biopsies in the same slice with identical mutations; in cases where more than two biopsies are positive, the path of the clone has been arbitrarily drawn. Solid grey regions represent biopsies that were not sequenced due to quality control issues. Gridded grey regions represent non-tubular regions of tissue.

**Supplementary Figure 3 – Individual gene plots showing the location of spontaneous mutations identified in testicular biopsies for AKT3 (A), APC (B), BRAF (C), CBL (D), FGFR3 (E), KRAS (F), LRP5 (G), MAP2K1 (H), MAP2K2 (I), NF1 (J), RAF1 (K), RET (L), and SOS1 (M).** (Panel I) Validated variants (with VAF on *y*-axis) positioned along the amino acid sequence of the relevant protein (*x*-axis, see Panel V). (Panel II) Location and size of amplicons used to sequence main hotspots of the relevant genes are plotted on the *x*-axis. Median coverage per amplicon is plotted on the *y*-axis. Line indicates coverage cut-off of 5,000x. (Panel III) Number of reported constitutional variants encoding amino acid substitutions associated with developmental disorders (sqrt scale). (Panel IV) Number of reported somatic amino acid substitutions in cancer (COSMIC v82). (Panel V) Protein domains. Annotations are based on the transcripts accessions listed in the methods.

**Supplementary Figure 4 - Variant allele frequencies of *KRAS* c.182A>G (p.Gln61Arg) and *LRP5* c.291C>T (p.Ala97Ala) in all 288 samples.**

**Supplementary Figure 5 – Heatmap of *NF1* c.2280G>A and *KRAS* c.35G>A.** Heatmap of G>A variants in *NF1* (called in 9 biopsies in Tes4 – surrounded by black lines) and *KRAS* (called in 6 biopsies in Tes4 – surrounded by black lines) reveals that there were a number of additional pieces with relatively high levels of the NF1 c.2280G>A variant that were not called. Heatmaps of the same variants in Tes1 and Tes2 demonstrate that the higher levels are specific to Tes4.

**Supplementary Figure 6 – Mutation loadings per sample.** Note that a number of samples show excessive C>A(G>T) mutations, which is typically associated with oxidative stress during the experimental procedure. Filtering of specific sample-mutation combinations and amplicons with excessive number of variants resulted in 6054 variant calls.

**List of Supplementary Tables and other supplementary files:**

**Supplementary Table 1 – Literature review showing loci with evidence for selfish selection**

**Supplementary Table 2 – Coverage analysis of 500 amplicons**

**Supplementary Table 3 – Table of prioritized calls (Tiers 1, 2, 3, 4)**

**Supplementary Table 4 – Variant calls in replicate samples**

**Supplementary Table 5 – Sample information**

**Supplementary Table 6 – Primers and smMIPs**

